# A sleep-regulatory circuit integrating circadian, homeostatic and environmental information in *Drosophila*

**DOI:** 10.1101/250829

**Authors:** Angélique Lamaze, Patrick Krätschmer, James E. C. Jepson

## Abstract

In the wild, when to go to sleep is a critical decision. Sleep onset is controlled by two processes: the circadian clock, and a homeostat measuring sleep drive [1, 2]. Environmental stimuli must also clearly intersect with the circadian clock and/or homeostat so that sleep is initiated only when appropriate. Yet how circadian, homeostatic and environmental cues are integrated at the circuit level is unclear. Recently, we found that DN1p clock neurons in *Drosophila* act to prolong morning wakefulness at elevated ambient temperatures [3]. Here we show that a subset of DN1p neurons exhibit temperature-sensitive increases in excitability, and define an output pathway linking DN1p neurons to downstream sleep-regulatory circuits. We show that DN1p neurons project axons to a subdomain of the Anterior Optic Tubercle (AOTU), and here make inhibitory synaptic connections with sleep-promoting tubercular-bulbar (TuBu) neurons. Using unbiased trans-synaptic labeling, we show that these TuBu neurons form synaptic connections with R-neurons innervating the ellipsoid body, subsets of which control homeostatic sleep drive [4]. DN1p excitability is clock-dependent, peaking in the late night and early morning [5]. Thus, integration of circadian and thermo-sensory information by DN1p neurons and subsequent inhibition of sleep-promoting TuBu neurons provides a mechanism by which an environmental stimulus can regulate sleep onset during a specific compartment of the day-night cycle. Furthermore, our results suggest that the AOTU functionally links circadian and sleep homeostat circuits in *Drosophila*.

## RESULTS AND DISCUSSION

### Consolidated sleep onset in *Drosophila* is gated by circadian and thermal cues

Using the standard definition of a *Drosophila* sleep bout as a 5 min period of inactivity (measured using the *Drosophila* Activity Monitor (DAM) system [6]), we recently showed that elevating ambient temperature from 22°C to ≥ 30°C delays onset of the first morning sleep bout in *Drosophila* males [3]. However, *Drosophila* sleep is often initially fragmented during the early morning at low-to-medium temperatures (Figure 1A). To gain a more robust view of how temperature changes affect consolidated sleep, we generated an R-based program capable of quantifying up to 30 distinct sleep parameters using raw DAM system data (see STAR Methods). We used this program to analyse how various properties of sleep architecture were modulated by temperature increases (Figure S1A-J), particularly the daytime longest sleep bout (dLSB).

**Figure 1:**
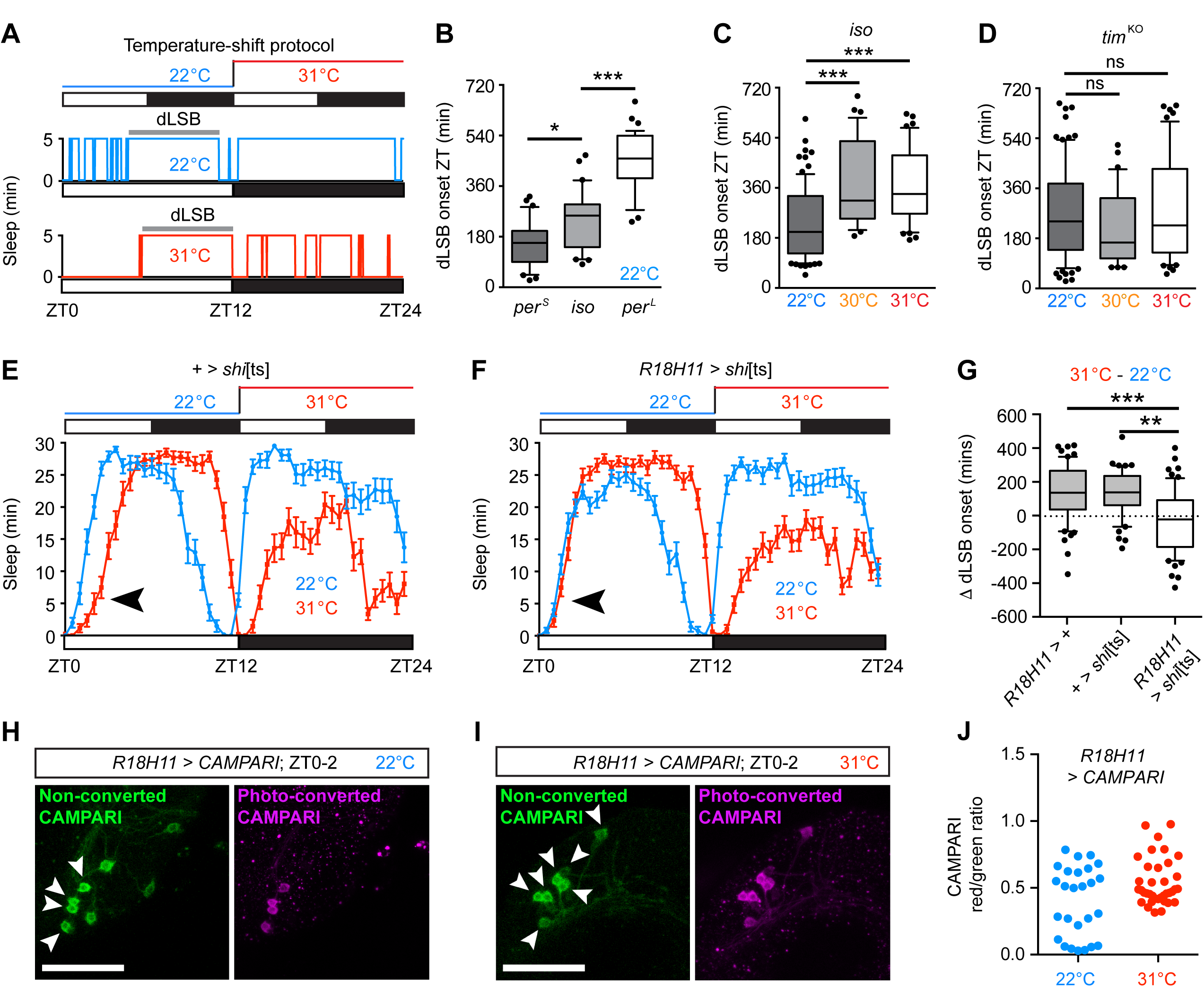
Thermo-sensitive DN1p neurons are required for delaying sleep onset at elevated temperature. (A) Upper panel: protocol used for temperature-shift experiments. Flies were housed under 12 h light: 12 h dark (12L: 12D) conditions. Sleep was measured on two consecutive days at ambient temperatures of 22°C and 31°C respectively. Light bars: day. Dark bars: night. Lower panels: sleep bouts for a single male fly under the above conditions. Note the fragmented sleep early in the morning at 22°C, before a longer consolidated sleep bout (grey bar). dLSB: longest sleep bout. At 31°C, morning wakefulness is enhanced, and onset of the dLSB is delayed. (B) Onset of the dLSB in *iso* control males, *per*^S^ and *per*^L^ mutants at 22°C. In these and all subsequent Tukey box plots, boxes show the 25^th^, median and 75^th^ percentiles. Whiskers show 1.5x the interquartile range. Dots represent outliers. *per*^S^: n = 30; *iso*: n = 31; *per*^L^: n = 37. (C) Onset of the dLSB in *iso* controls at different ambient temperatures. 22°C: n = 82; 30°C: n = 35; 31°C: n = 47. (D) Onset of the dLSB in *timeless* knockout (*tim*^KO^) mutants. 22°C: n = 85; 30°C: n = 33; 31°C: n = 52. (E) Overlaid mean sleep levels across two consecutive 24 h periods at 22°C and 31°C in control adult male flies harbouring the UAS-*shi*[ts] transgene alone (+ > *shi*[ts]). Note the delay in the onset of sleep during the day at elevated temperature (arrow). Error bars represent SEM. (F) Mean sleep levels in adult male flies following acute inhibition of DN1p synaptic output (*R18H11* > *shi*[ts]) at 31°C compared to the previous day at 22°C where DN1p synaptic output was uninhibited. Note the absence of a delay in morning sleep onset at 31°C (arrow). (G) Box plots showing median difference in onset of the dLSB following shifts from 22°C to 31°C in control and experimental male flies. *R18H11* > +: n = 63; + > *shi*[ts]: n = 50; *R18H11* > *shi*[ts]: n = 63. (H, I) Confocal images showing non-converted and photo-converted CAMPARI in *R18H11*-positive DN1p neurons at ZT0-2 at either 22°C (H) or 31°C (I). Arrows point to *R18H11*-positive DN1p neurons where robust photo-conversion, and thus elevated neuronal excitability, can be detected. Scale bars, 50 μm. (J) Dot plot illustrating ratios of non-converted to photo-converted CAMPARI in *R18H11*-positive DN1p neurons at either 22°C or 31°C. At 22°C, two sub-populations are apparent (minimal or robust photo-conversion), whereas at 31°C DN1p neurons show a higher and more uniform degree of CAMPARI photo-conversion. 22°C: n = 26 neurons from 4 brains. 31°C: n = 37 neurons from 7 brains. *p < 0.05, **p < 0.01, ***p < 0.001, ns – not significant, Kruskal Wallis test with Dun’s post-hoc test. See also Figure S1.

Given the prolonged inactivity during the dLSB (Figure 1A), we reasoned that this parameter would yield a more accurate representation of when consolidated sleep was occurring. We measured the onset and offset of the dLSB in control flies and in mutants with short and long circadian periods (*per*^S^ and *per*^L^) [7]. We observed an advance in the onset of the dLSB in *per*^S^ mutants and a delay in *per*^L^ relative to control flies at 22°C (Figure 1B) while offset of the dLSB was similarly advanced in *per*^S^ mutants (Figure S1A), demonstrating that timing of the dLSB is clock-dependent. The timing of the dLSB was also temperature-sensitive. Shifting flies from 22°C to ≥ 30°C resulted in a significant delay in both the onset and offset of the dLSB (Figures 1A, C and Figure S1B). Importantly, delayed onset at ≥ 30°C was suppressed in male flies lacking a functional circadian clock (*tim*^KO^) (Figure 1D) [3]. Thus, circadian and thermo-sensory pathways interact to modulate onset of the dLSB.

### Thermo-sensitive DN1p neurons delay morning sleep

Which clock neurons influence delay of the dLSB at high temperature? DN1p clock neurons promote delay of the first morning sleep bout at high temperatures [3]. Thus, we tested whether synaptic output from DN1p neurons was also required to delay the dLSB at 31°C. To suppress synaptic output from DN1p neurons, we expressed a temperature-sensitive inhibitor of endocytosis (*shi*[ts]) using the *R18H11*-Gal4 driver, which labels ~ 7-8 DN1p neurons [8, 9]. Expression of *shi*[ts] blocks synaptic release at 31°C but not 22°C via inhibition of synaptic vesicle recycling, facilitating acute inhibition of DN1p output at high temperature. As expected, control flies exhibited a delay in the onset of the dLSB at 31°C compared to 22°C (Figure 1E, G). In contrast, inhibiting synaptic release from *R18H11*-positive DN1p neurons blocked this effect (Figure 1F, G). Both the delay in onset and reduction in length of the longest night sleep bout at 31°C relative to 22°C was unaffected by DN1p inhibition (Figure 1E, F and Figure S1K-M). DN1p neurons therefore modulate the timing of consolidated sleep during the day but not the night in response to temperature changes.

We next asked whether the excitability of DN1p neurons was temperature-sensitive. To examine this, we expressed CAMPARI in DN1p neurons. When stimulated with UV light, the CAMPARI fluorophore undergoes green-to-red photo-conversion in high intracellular calcium, yielding an optical read-out of neuronal activity [10]. We utilised the relatively calcium-insensitive V398D variant to increase the dynamic range of CAMPARI [10], and performed photo-conversion experiments at Zeitgeber Time (ZT0-2) either at 22°C or following a shift from 22°C to 31°C at ZT0. Following photo-conversion, we measured the ratio of non- and photo-converted CAMPARI fluorescence in DN1p cell bodies (Figure 1H-J). At ZT0-2 and 22°C, approximately half of *R18H11*-DN1p neurons exhibited robust levels of photo-converted CAMPARI, indicative of high neuronal excitability (Figure 1H, J). In contrast, at the same time-point but at 31°C, we detected a more uniform increase in CAMPARI photo-conversion (Figure 1I, J). Indeed, the coefficient of variation of the ratio of photo-converted to non-converted CAMPARI was higher at 22°C relative to 31°C (0.65 vs. 0.32), and the population variance at 22°C was also significantly higher relative to 31°C (p = 0.026, F-test). These results suggest that DN1p clock cells contain a thermo-sensitive sub-population that act in a wake-promoting circuit in the morning.

### DN1p neurons form synaptic connections in the Anterior Optic Tubercle

To identify relevant circuits downstream of DN1p neurons, we co-expressed distinct fluorophores localised to presynaptic and dendritic domains (UAS-*syt-GFP* and UAS-*DenMark* respectively) using *R18H11*-Gal4 (Figure 2A). Confocal imaging revealed DN1p dendrites in the dorsal posterior protocerebrum, within the region in which thermo-sensory TrpA1-expressing neurons form synaptic contacts with DN1p neurons [3]. DN1p presynaptic boutons innervated two neuropil regions: the pars intercerebralis [11], but also the Anterior Optic Tubercle (AOTU) (Figure 2A). Expression of *syt*-GFP using a distinct driver that also labels DN1p neurons (*clk4*.1M-Gal4) confirmed the presence of DN1p synapses in this region (Figure S2A) [12, 13].

**Figure 2.**
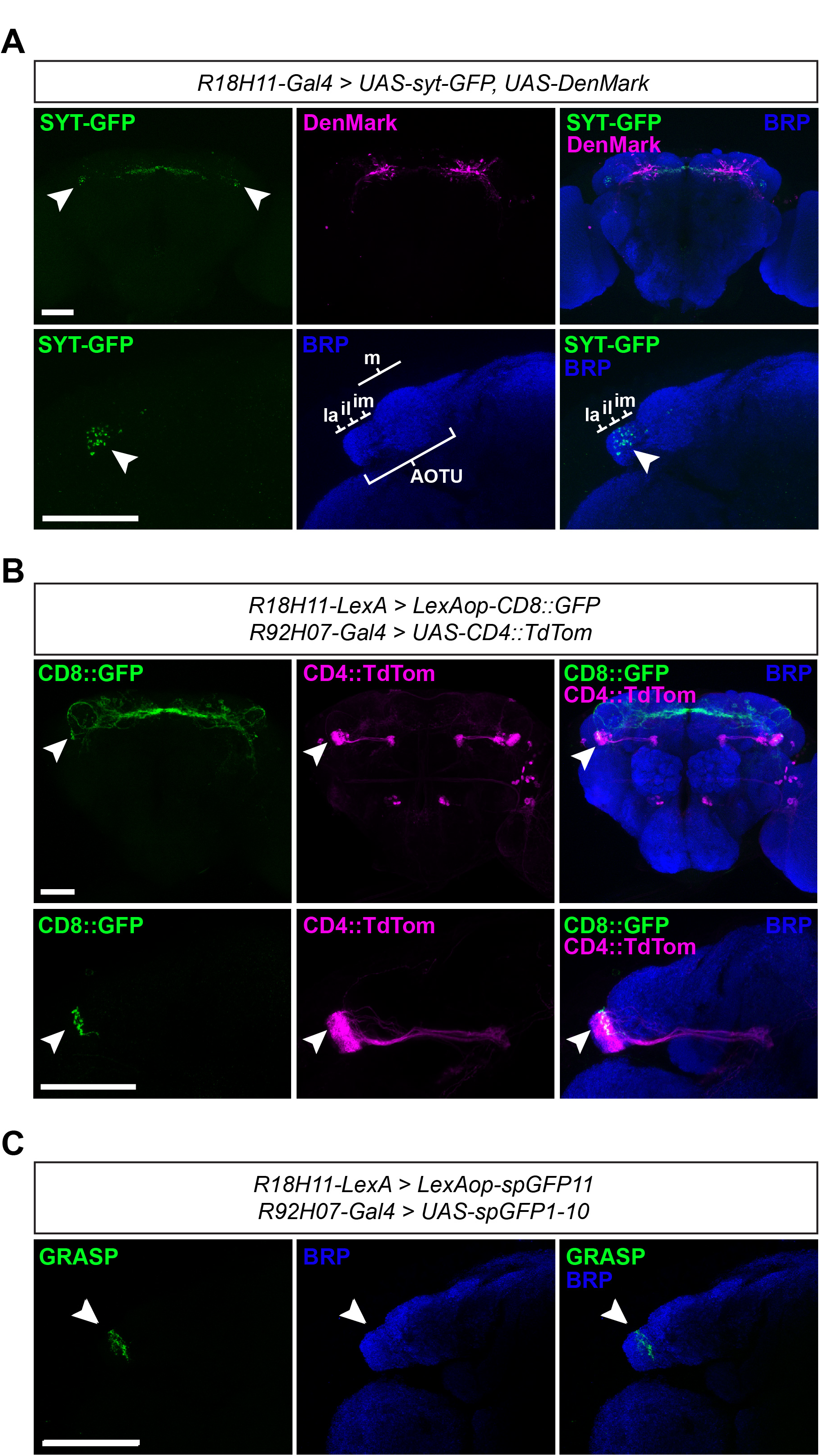
DN1p neurons form synaptic contacts with TuBu neurons in
the AOTU. (A) Upper panel: confocal images showing co-expression of presynaptic and dendritic markers in adult male DN1p neurons. Arrows: pre-synaptic SYT-GFP puncta in the anterior optic tubercle (AOTU). Lower panel: high-resolution image of DN1p presynaptic domains (arrow) in the AOTU. Subdomains of the AOTU are as follows. La: lateral, il: intermediate lateral, im: intermediate medial, m: medial [14]. BRP: Bruchpilot, a presynaptic neuropil marker.
(B) Upper panel. Confocal images showing projection patterns of DN1p neurons (labelled using *R18H11*-LexA) and *R92H07*-Gal4-positive TuBu neurons. Distinct membrane-tagged fluorophores were expressed in each cell type. Arrows point to the AOTU region. Lower panel: high-resolution image of the AOTU showing overlap (arrows) between DN1p axons/synapses and dendritic regions of *R92H07*- TuBu neurons.
(C) High-resolution confocal image showing synaptic connectivity between DN1p and *R92H07-* TuBu neurons, as demonstrated through GRASP. Arrows point to reconstituted GFP signal formed in the AOTUil following expression of membrane-tagged GFP fragments in DN1p and *R92H07-*TuBu neurons. Scale bars: 50 μm. See also Figure S2.

The AOTU is a component of the Anterior Visual Pathway, linking neurons in the optic lobe medulla to the Ellipsoid Body (EB) [14], a domain of the central complex involved in motor control [15], navigational learning and orientation [16, 17], and importantly, homeostatic sleep drive [4]. Dendrites from tubercular-bulbar (TuBu) neurons innervate the AOTU and project axons to the bulb (also known as the lateral triangle), where they synapse onto EB ring (R-) neurons [14, 18]. The AOTU can be divided into distinct regions: the medial (AOTUm), intermediate medial (AOTUim), intermediate lateral (AOTUil) and lateral (AOTUla) (as defined in ref. [14]) (Figure 2A). DN1p synapses predominantly localise to the AOTUil, with a small proportion innervating the AOTUim and AOTUla (Figure 2A). We identified TuBu neurons innervating the AOTUil that can be labelled using the *R92H07*-Gal4 driver (Figure 2B). To confirm synaptic connectivity between DN1p and *R92H07*-TuBu neurons, we first used orthogonal LexA- and Gal4-drivers to express different fluorophores in each cell type, revealing close association of DN1p axons/synapses with *R92H07*-TuBu dendrites (Figure 2B). Using the same approach we next performed GRASP experiments by expressing complementary fragments of GFP in DN1p and *R92H07*-TuBu neurons [19]. Confocal imaging revealed reconstituted GFP fluorescence specifically in the AOTUil region (Figure 2C and Figure S2B), suggesting synaptic connectivity between DN1p and TuBu neurons.

We also tested whether DN1p neurons formed connections with TuBu neurons in the more lateral region of the AOTU (AOTUla). Using the *R83H09*-Gal4 driver we labelled AOTUla TuBu neurons (Figure S2E). DN1p synapses/axons tiled the boundary of the dendritic domain formed by *R83H09*-Tubu neurons (Figure S2C). However, in contrast to *R92H07*-Tubu neurons in the AOTUil, no GRASP signal between DN1p and *R83H09*-Tubu neurons in the AOTUla was observed (Figure S2D). Thus, DN1p neurons appear to physically associate predominantly with TuBu neurons in the AOTUil region.

### DN1p neurons inhibit TuBu neurons

We next examined whether DN1p and *R92H07*-TuBu neurons are functionally connected. The excitability of DN1p neurons oscillates in a clock-dependent manner, with a peak in the late night/early morning and a trough in the late day/early night [5]. Thus, we wondered whether the excitability of *R92H07*-TuBu neurons also oscillated, and if so, whether such oscillation was positively or negatively correlated with DN1p neurons. To do so, we again utilised CAMPARI [10]. Consistent with previous patch-clamp analysis [5], excitability of DN1p neurons at 31°C was higher in the morning (a period of increased wakefulness) relative to the afternoon when male flies are largely asleep (Figure 3A). In *R92H07*-TuBu neurons the opposite was the case (Figure 3B). This suggests that DN1p neurons inhibit *R92H07*-TuBu neurons. To test this, we expressed ChannelRhodopsin2-XXL (ChR2-XXL) in DN1p neurons and the GCamP6s reporter of intracellular calcium in *R92H07*-TuBu neurons [20, 21]. We optogenetically activated DN1p neurons at ZT9 and 31°C – a time-point and temperature at which excitability of DN1p neurons is low and *R92H07*-TuBu neurons high (Figure 3A, B) [5] – and measured changes in intracellular calcium in *R92H07*-TuBu neurons. In control brains expressing GCamP6s in *R92H07*-TuBu neurons, excitation with UV light causes an increase in GCamP6s fluorescence due to an overlap with the GCamP6s excitation spectra (Figure 3C). However, when UV light-activated ChR-XXL was simultaneously expressed in DN1p neurons, this increase in GCamP6s fluorescence was greatly reduced, indicative of a parallel reduction in intracellular calcium (Figure 3C). Collectively, the above results indicate that DN1p neurons inhibit *R92H07*-TuBu neurons.

**Figure 3.**
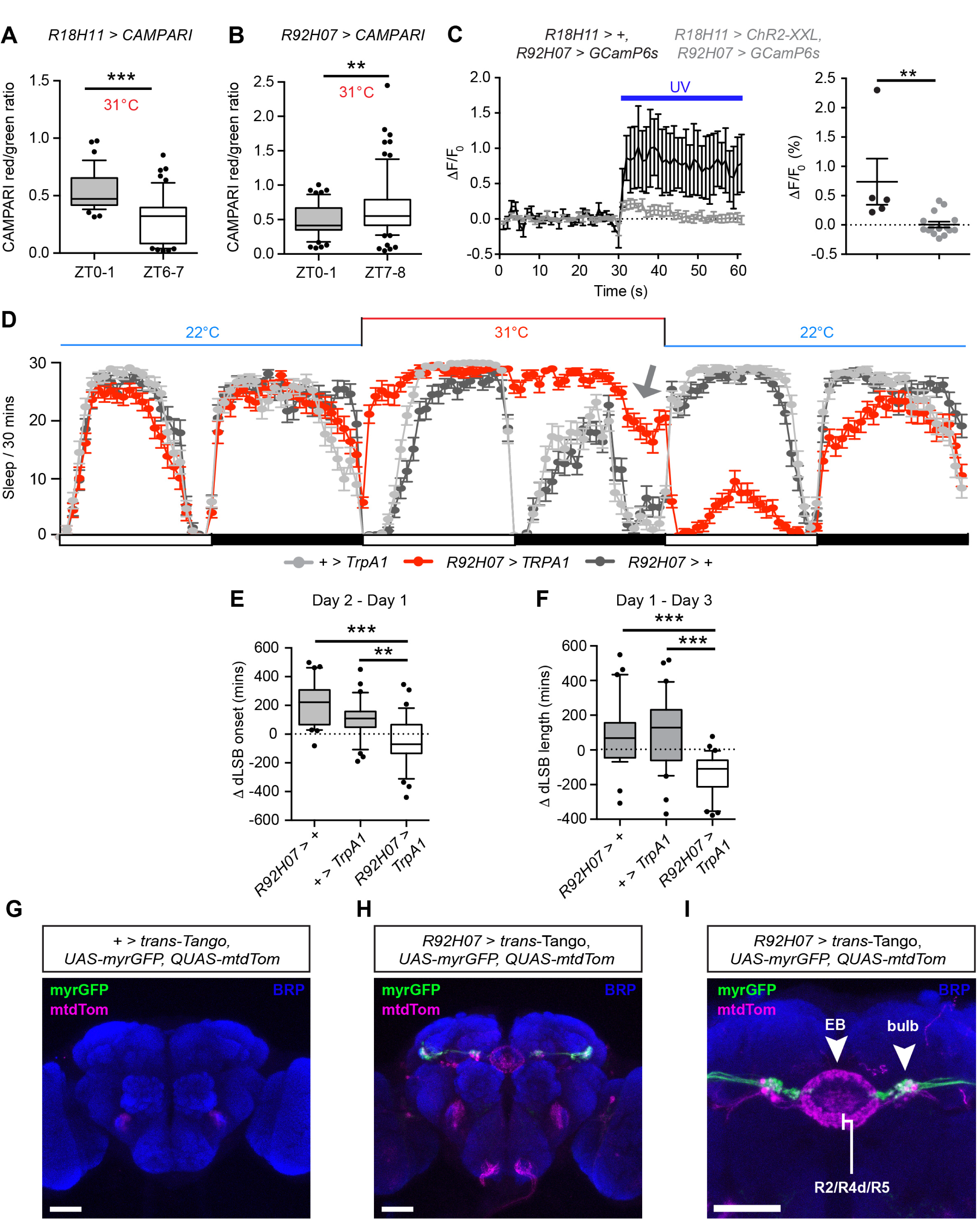
DN1p neurons inhibit sleep-promoting TuBu neurons. (A-B) Box plots showing ratios of photo-converted (red) and non-converted (green) CAMPARI fluorescence at the stated time-points in *R18H11*-DN1p (A) and *R92H07-*TuBu neurons (B). *R18H11*-DN1p neurons (A): ZT0-1, n = 37 neurons from 7 brains; ZT7-8, n = 52 neurons from 8 brains. *R92H07-*TuBu neurons (B): ZT0-1, n = 51 neurons from 7 brains; ZT6-7, n = 75 neurons from 8 brains. (C) Left panel: mean change in GCamP6s fluorescence in *R92H07*-TuBu neurons (*R92H07*-LexA > LexAop-*GCamP6s*) following UV light activation of ChR2-XXL in DN1p neurons (grey; *R18H11*-Gal4 > UAS-*ChR2-XXL*), or solely due to UV light stimulation (black). Error bars represent SEM. Right panel: dot plots of individual cellular changes in GCamP6s fluorescence following UV light stimulation in the presence (grey) or absence (black) of ChR2-XXL expressed in DN1p neurons. Mean and SEM are shown. (D) Overlaid mean sleep levels across three consecutive 24 h periods at 22°C (control day), 31°C (TrpA1 activation) and 22°C (recovery day), in the stated genotypes. Arrow: reduced sleep prior to lights-on during *R92H07*-TuBu activation. Error bars represent SEM. (E, F) Box plots showing median difference in the onset of the dLSB following shifts from 22°C to 31°C (E) or in the length of the dLSB between control and recovery days (F) in control and experimental male flies. *R92H07* > +: n = 29; + > *TrpA1*: n = 34; *R92H07* > *TrpA1*: n = 36. **p < 0.01, ***p < 0.001, Mann-Whitney U-test (A-C) or Kruskal Wallis test with Dun’s post-hoc test (E, F). (G) Confocal image of a control *trans*-Tango male adult brain lacking any Gal4 driver. BRP, Bruchpilot. (H, I) Confocal images of *trans*-Tango driven by *R92H07*-Gal4. myr-GFP marks *R92H07*-TuBu neurons. Projections from a small population of non-TuBu neurons are also labelled at very low levels. mtdTomato (mtdTom) labels post-synaptic neurons. Note overlap of myr-GFP and mtdTomato signals in the superior bulb (arrow), with projections from downstream neurons innervating the ellipsoid body rings (EB, arrow) (I). EB rings innervated by R2-/R4d-/R5-neurons are shown (I). Scale bars: 50 μm. See also Figure S3.

### TuBu neurons in the AOTUil are sleep-promoting

DN1p neurons are wake-promoting during the morning at high temperatures [3] (Figure 1F). Thus, if DN1p neurons inhibit *R92H07*-TuBu neurons, *R92H07*-TuBu neurons should promote sleep. To test this, we expressed TrpA1 (a temperature-gated cation channel) in *R92H07*-neurons and measured sleep following a shift from 22°C (a non-activating temperature) to 31°C (an activating temperature) [22]. Excitation of *R92H07*-neurons immediately and profoundly induced sleep throughout the day, whereas control flies exhibit prolonged morning wakefulness (Figure 3D, E). Sleep loss during the night caused by elevated temperature was also suppressed by *R92H07*-TuBu neurons activation (Figure 3D). Note that at the end of the night, morning anticipation (a clock-driven locomotor behavior) remained intact following *R92H07*-TuBu excitation (Figure 3D, grey arrow). In addition, the total number of sleep bouts during the night was not significantly reduced (Figure S3A, B). Thus, these flies are able to exhibit waking behavior and are not simply paralyzed. On the following recovery day at 22°C, flies in which sleep was previously initiated via *R92H07*-TuBu excitation exhibited a strong ‘negative rebound’, showing a reduction in the length of the dLSB relative to the initial control day at 22°C, in contrast to an increase in controls (Figure 3D, F). These results suggest that *R92H07*-TuBu neurons are coupled to circuits that control homeostatic sleep drive, in contrast to other sleep-modulatory cell-types in the fly brain such as octopaminergic neurons [23]. To test whether suppressing *R92H07*-TuBu output impacted sleep at high temperature, we acutely blocked synaptic release from *R92H07*-TuBu neurons using *shi*[ts] (Figure S3C). Inhibition of *R92H07*-TuBu neurons reduced the length of the longest sleep bout during the day but not the night (Figure S3D-I), consistent with a role for *R92H07*-TuBu neurons in promoting day sleep. We note, however, that the effect of inhibiting *R92H07*-TuBu neurons on overall sleep architecture was subtle compared to acute activation (Figure S3E, Figure 3D). These results support a model in which additional sleep-promoting TuBu neurons within the AOTUil are unmarked by the *R92H07-*Gal4 driver and are still available to promote sleep during inhibition of *R92H07*-TuBu neurons.

### Sleep-regulatory TuBu neurons form synaptic connections with diverse ellipsoid body R-neurons

Finally, we sought to identify circuits downstream of *R92H07*-TuBu neurons. To do so, we utilised a recently devised system for unbiased trans-synaptic labelling: *trans*-Tango [24]. This method involves expression of a signalling molecule (Glucagon) tethered to presynaptic domains of a neuron of interest. In parallel, a post-synaptic Glucagon receptor is expressed in all neurons, and is connected to a signalling pathway yielding transcription of the fluorophore mtdTomato following binding of Glucagon to its receptor on post-synaptic neurons (see ref. 24 for details). Presynaptic neurons are simultaneously labelled by expressing a distinct fluorophore (myr-GFP) [24]. We applied this method to *R92H07*-TuBu neurons. Control flies containing *trans*-Tango but no driver exhibited non-specific mtdTomato fluorescence in the antennal mechanosensory and motor centre (Figure 3G). In contrast, when *trans*-Tango was driven by *R92H07*-Gal4, we observed overlapping myr-GFP and mtdTomato fluorescence in the superior region of the bulb (Figure 3H, I), as indicated by the dorsal location of myr-GFP and mtdTomato relative to the EB midline (Figure 3I). Neurons post-synaptic to *R92H07*-TuBu neurons projected to the EB (Figure 3I, J). From the location of their dendrites in the superior bulb and axonal projections in the outer segments of the EB, these may represent R2-, R4d- and R5 neurons (Figure 3I) [14, 17, 18]. R1/R3 neurons in the central domain of the EB were also visible, albeit more weakly (Figure 3I). However, since the dendrites of R1/R3 neurons innervate the inferior bulb [14, 18], these are unlikely to be functionally connected to *R92H07*-TuBu neurons, which project to the superior bulb (Figure 3I). In summary, our results define a tripartite sleep-regulatory circuit linking DN1p neurons to the EB via the AOTU.

## Conclusions

Here we elucidate a circuit mechanism by which sleep during the early morning is suppressed at elevated temperatures. We propose that circadian and thermo-sensory information is concurrently encoded in the excitability of DN1p clock neurons. Subsequently, sleep-regulatory information is transmitted from DN1p neurons to the EB, a motor control centre [15, 25], via TuBu neurons innervating the AOTUil. This results in the suppression of locomotion resulting from sensory input such as visual stimuli and initiation of sleep [4, 17, 26]. Our trans-synaptic labelling experiments linking TuBu to EB R-neurons are consistent with recent studies defining these two cell-types as neighboring modules of the anterior visual pathway [14], [18]. Importantly, R2-neurons also receive recurrent feedback from sleep-promoting neurons in the dorsal fan-shaped body [26], and enhance sleep drive when artificially activated [4]. R2-neurons have therefore been proposed to act as a component of the sleep homeostat [4]. *R92H07*-TuBu neurons form synaptic contacts with R-neurons innervating multiple EB rings (Figure 3I). Thus, the sleep-promoting action of *R92H07*-TuBu neurons is likely to occur via simultaneous modulation of multiple subsets of EB neurons, the net effect of which is to suppress locomotion and promote sleep. The inhibition of *R92H07*-TuBu neurons by DN1p neurons provides a mechanism to promote wakefulness when DN1p activity is high: during the early morning and at high temperatures (Figures 1I and 3A) [3, 5]. In concert with previous data [3, 4], our results suggest that the DN1p > TuBu > R-neuron circuit represents a physical link between clock, sensory, and sleep homeostat neurons, enabling circadian and environmental cues to dynamically regulate sleep onset.

## ACKNOWLEDGEMENTS

We thank Gilad Barnea, Kyunghee Koh, Francois Rouyer and Ralf Stanewsky for *Drosophila* stocks; and Ko-Fan Chen, Simon Lowe, Kyunghee Koh and Francois Rouyer for helpful comments on the manuscript. Ko-Fan Chen provided technical input relating to live-imaging and driver lines labelling the AOTU. This study was supported by a UCL Start-Up Fund to J.E.C.J.

## AUTHOR CONTRIBUTIONS

Conceptualization: A.L and J.E.C.J. Methodology: A.L. Software: P.K.

Validation: A.L. Formal Analysis: A.L and P.K. Investigation: A.L. Writing –

Original Draft: J.E.C.J and A.L. Writing – Review and Editing: A.L, P.K and

J.E.C.J. Visualisation: A.L, P.K and J.E.C.J. Supervision: J.E.C.J. Project

Administration: J.E.C.J. Funding Acquisition: J.E.C.J.

## DECLARATION OF INTERESTS

The authors declare no competing interests.

## STAR METHODS

### CONTACT FOR REAGENT AND RESOURCE SHARING

Further information and requests for resources and reagents should be directed to and will be fulfilled by the Lead Contact, James Jepson

### EXPERIMENTAL MODEL AND SUBJECT DETAILS

Fly strains and crosses were reared on standard yeast-containing fly flood at a constant temperature of 25°C, housed under 12 h: 12 h light-dark cycles (LD). *Drosophila* lines used for behavioral analysis were outcrossed for a minimum of four generations into an isogenic (iso31) background. Individual 2-4 day old males were used in all behavioral experiments. *per*^L^ and *per*^S^ mutant lines were kind gifts from Francois Rouyer and Ralf Stanewsky. *Trans*-Tango lines were a kind gift from Gilad Barnea.

## METHOD DETAILS

### Behavioral assays

Individual males were loaded into glass tubes containing 2% agar and 4% sucrose. Sleep measurements were performed using the Drosophila Activity Monitor (DAM) system (Trikinetics, MA, USA) [6]. For all experiments shown in this manuscript, Trikinetics monitors were housed in temperature- and light-controlled incubators (LMS, UK) as described previously [3]. Sleep graphs derived from Trikinetics data were generated using GraphPad Prism 6.

### Immunohistochemistry

Adult male *Drosophila* brains were immuno-stained as described previously [27]. Brains were fixed in 4% paraformaldehyde at RT for 20 min, and blocked in 5% goat serum at RT for 1 h. Primary antibodies used were as follows: rabbit anti-DsRed (Clontech) – 1:2000; mouse anti-Bruchpilot (nc82, Developmental Studies Hybridoma Bank (DSHB)) – 1:200; chicken anti-GFP (ThermoFisher) – 1:1000. Alexa-fluor secondary antibodies (goat anti-rabbit 555, goat anti-chicken 488 and goat anti-mouse 647; ThermoFisher) were used at 1:2000 except for labelling anti-BRP where goat anti-mouse 647 at a dilution of 1:500 was used.

### Confocal imaging and optogenetics

All confocal and optical imaging experiments were performed using an inverted Zeiss LSM 710 confocal microscope. For experiments involving CAMPARI, crosses were performed at 25°C. Subsequently, 1-3 days old flies were entrained in LD 12-12 cycles at 22°C. 48 h prior to the experiment, adult male *Drosophila* were transferred to individual glass tubes to facilitate catching without CO_2_ or cold-induced anaesthesia. On the experimental day, individual flies were manually caught and brains were subsequently dissected in HL3.1 buffer, of composition: 70 mM NaCl, 5 mM KCl, 20 mM MgCl_2_, 1.5 mM CaCl_2_, 10 mM NaHCO_3_, 5 mM Trehalose, 115mM sucrose, 5mM HEPES, pH 7.2. UV exposure was undertaken using a high power mercury lamp for 2 min. When testing for changes in CAMPARI photo-conversion following a rise in ambient temperature from 22°C to 31°C, the dissection buffer was pre-heated to 31°C in order to maintain the experimental temperature. For experiments involving GCamP6s, brains were dissected in the same dissection buffer as above. To excite GCamP6s, a 514 nm laser was used, whereas to activate ChR2XXL, a 405nm laser was applied.

## QUANTIFICATION AND STATISTICAL ANALYSIS

Since many of the datasets derived from sleep experiments exhibited a non-normal distribution, the following statistical tests were used. For single comparisons, Mann-Whitney U-tests were used. When comparing multiple genotypes, Kruskal-Wallis tests were used, followed by Dunn’s post-hoc tests. All statistical analyses were performed using GraphPad Prism 6. For each dataset, details of statistical tests used, n-values, dispersion and precision measures can be found in the corresponding Figure Legends.

## DATA AND SOFTWARE AVAILABILITY

The custom-made R-based package for analysis of the duration, onset and offset of the longest sleep bout is detailed in, and can be downloaded from, the following Github link: https://github.com/PatrickKratsch/DAM_sleep_parameters

## Supplemental Information

**Figure S1.**
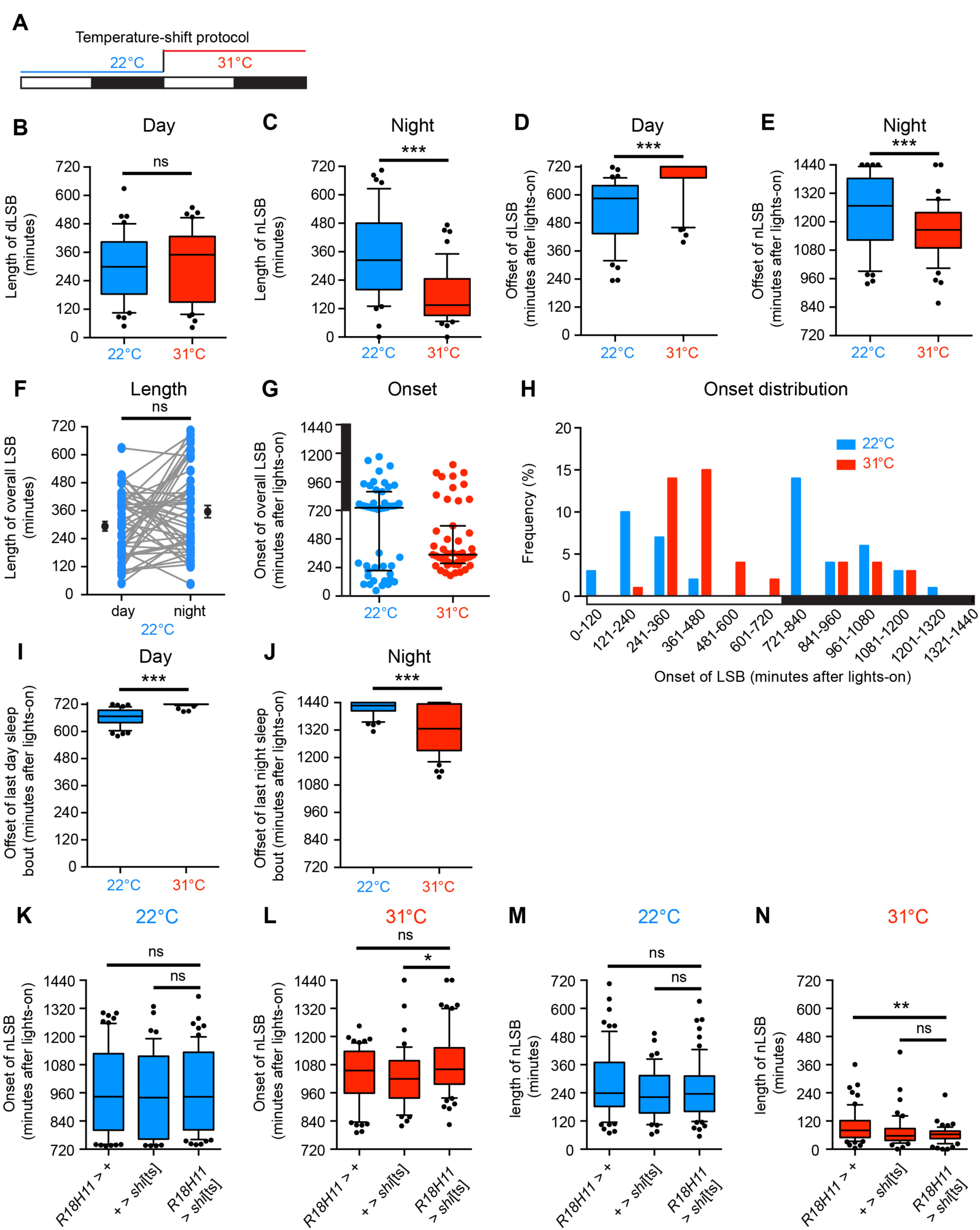
relating to Figure 1. Temperature-dependent changes in sleep architecture and role of DN1p clock neurons. (A) Paradigm used to test how a range of parameters describing sleep architecture change in response to elevated temperature. On day 1, adult male flies were housed at 22ºC, and on day 2, ambient temperature was raised to 31ºC. White bars: day. Black bars: night. Activity was recorded on both days using the DAM system and the custom-made R-program was used to quantify sleep parameters described in (B-J). All data presented in (B-J) are derived from n = 47 iso31 control adult male flies. (B-E) Changes in the length (B-C) and time of offset (D-E) of the longest sleep bouts (LSB) during the day (dLSB; B, D) or night (nLSB; C, E) in response to increased ambient temperature. Note that day and night sleep patterns respond differently to increased ambient temperature. The length of the daytime LSB is not changed (B), whereas the length of the night LSB is significantly reduced (C). Furthermore, the offsets of the day and night LSBs are also altered in an opposing manner, with offset of the day LSB being delayed (D), and offset of the night LSB advanced (E), consistent with a reduced length of night LSB. The distribution of data is illustrated using Tukey box plots, as described in Figure 1. (F) Paired comparison of the length of the day and night LSB in individual flies at 22ºC. Connecting bars indicate how these two parameters vary in individual flies. No significant difference in the average LSB between day and night was observed (black circle denotes mean ± SEM for day and night LSBs). This is consistent with *Drosophila* exhibiting a crepuscular pattern of rest/activity. (G, H) Distributions showing the onset of the LSB across the complete 24 h period (overall LSB), shown via dots plots (G) or histogram (H). White bars: day. Black bars: night. Median and interquartile ranges are shown. As described above in (F), at 22ºC the day and night LSBs are not significantly different in length. Correspondingly, for a population of flies housed at 22ºC, the overall LSB during 24 h can be initiated during either the day or the night, but is most commonly observed directly after lights-off. However, at 31ºC we observed that the overall LSB is more frequently initiated during the day, suggesting that at high temperatures flies shift from a crepuscular to a nocturnal pattern of activity. (I, J) Offset of the last day (I) and night (J) sleep bout at either 22ºC or 31ºC. Again, day and night sleep offset responds in an opposite manner to increased temperature, with the offset of the last day sleep bout being delayed, and the offset of the last night sleep bout advanced, at high ambient temperature. (K-N) Comparison of the onset (K, L) and length (M, N) of the night longest sleep bout (nLSB) at either 22ºC or 31ºC in adult male flies expressing the temperature-sensitive inhibitor of synaptic vesicle endocytosis (*R18H11* > *shi*[ts]) and controls. 22ºC (K, M): control condition in which dominant-negative temperature-sensitive *shibire* (*shi*[ts]) is inactive. 31ºC (L, N): DN1p inhibition by *shi*[ts] (*R18H11* > *shi*[ts]) and control flies examined under the same temperature conditions. *R18H11* > +: n = 63; + > *shi*[ts]: n = 50; *R18H11* > *shi*[ts]: n = 63. *p< 0.05, **p< 0.01, ***p< 0.001, ns – p > 0.05, Mann-Whitney U-test (B-F, I-J) or Kruskal-Wallis test with Dunn’s post-hoc test (K-N).

**Figure S2.**
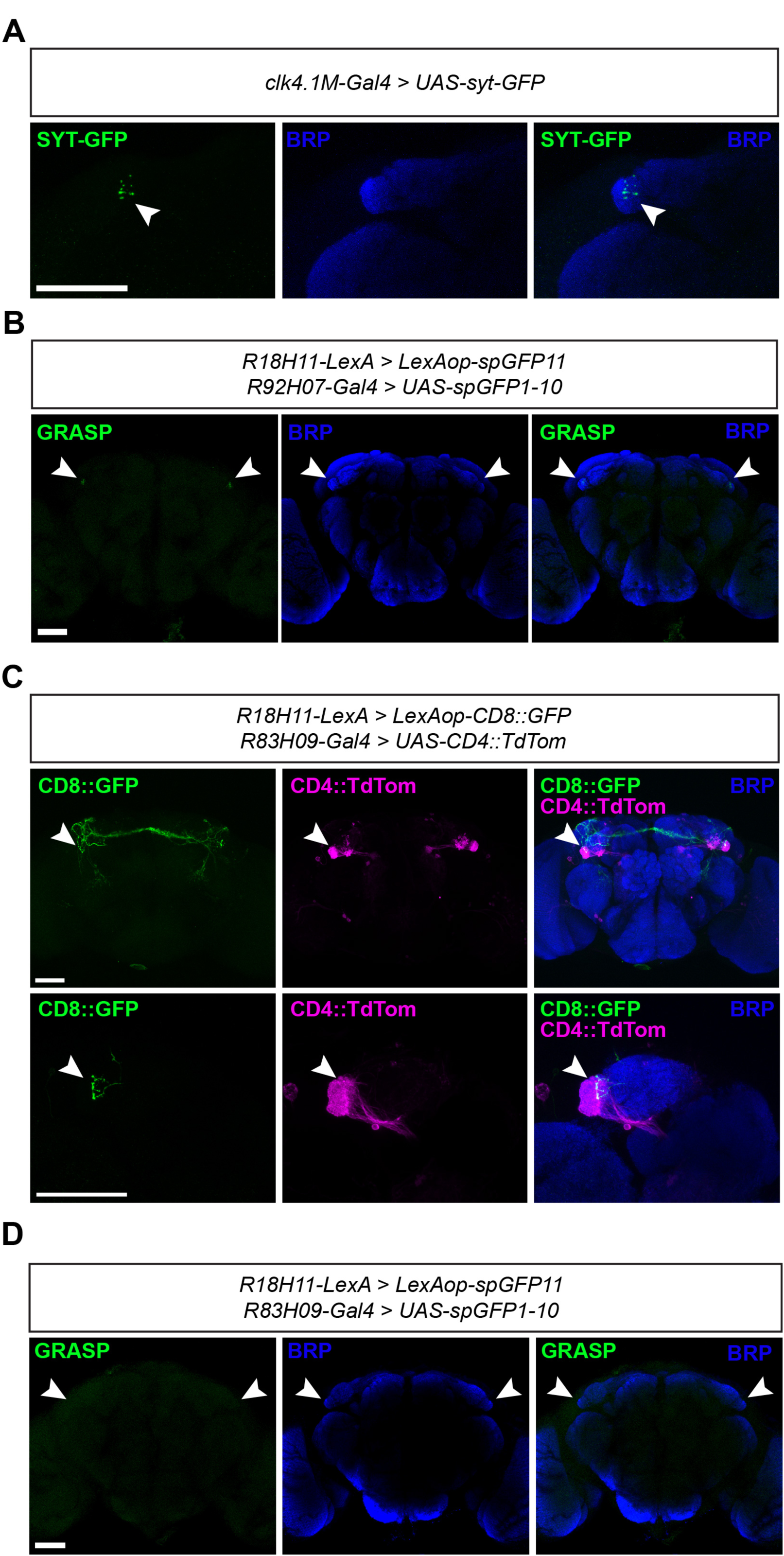
relating to Figure 2. DN1p neurons make synaptic connections with TuBu neurons in the AOTU. (A) Confocal images showing localisation of presynaptic SYT-GFP puncta derived from *clk4.1M*-Gal4-expressing adult male DN1p neurons. Arrows point to SYT-GFP puncta within the anterior optic tubercle (AOTU). BRP:Bruchpilot (a presynaptic neuropil marker). (B) Confocal images showing that GRASP signal obtained through expression of split-GFP fragments in DN1p neurons (via *R18H11*-LexA) and TuBu neurons (via *R92H07*-Gal4) is detected specifically in the AOTU (arrows), and not elsewhere in the central brain. (C) Dual labelling of DN1p and *R83H09*-positive TuBu neurons that innervate the lateral AOTU. Arrows point to DN1p presynaptic termini that tile the boundary of the lateral AOTU, and thus the dendritic domain of *R83H09*-TuBu neurons. (D) Confocal images showing lack of GRASP signal from split-GFP fragments expressed in DN1p neurons (via *R18H11*-LexA) and *R83H09*-TuBu neurons. Arrows point to the AOTU region. No GRASP signal was detected here or elsewhere in the *Drosophila* brain (compare with Figure S2A). All scale bars: 50 μm.

**Figure S3.**
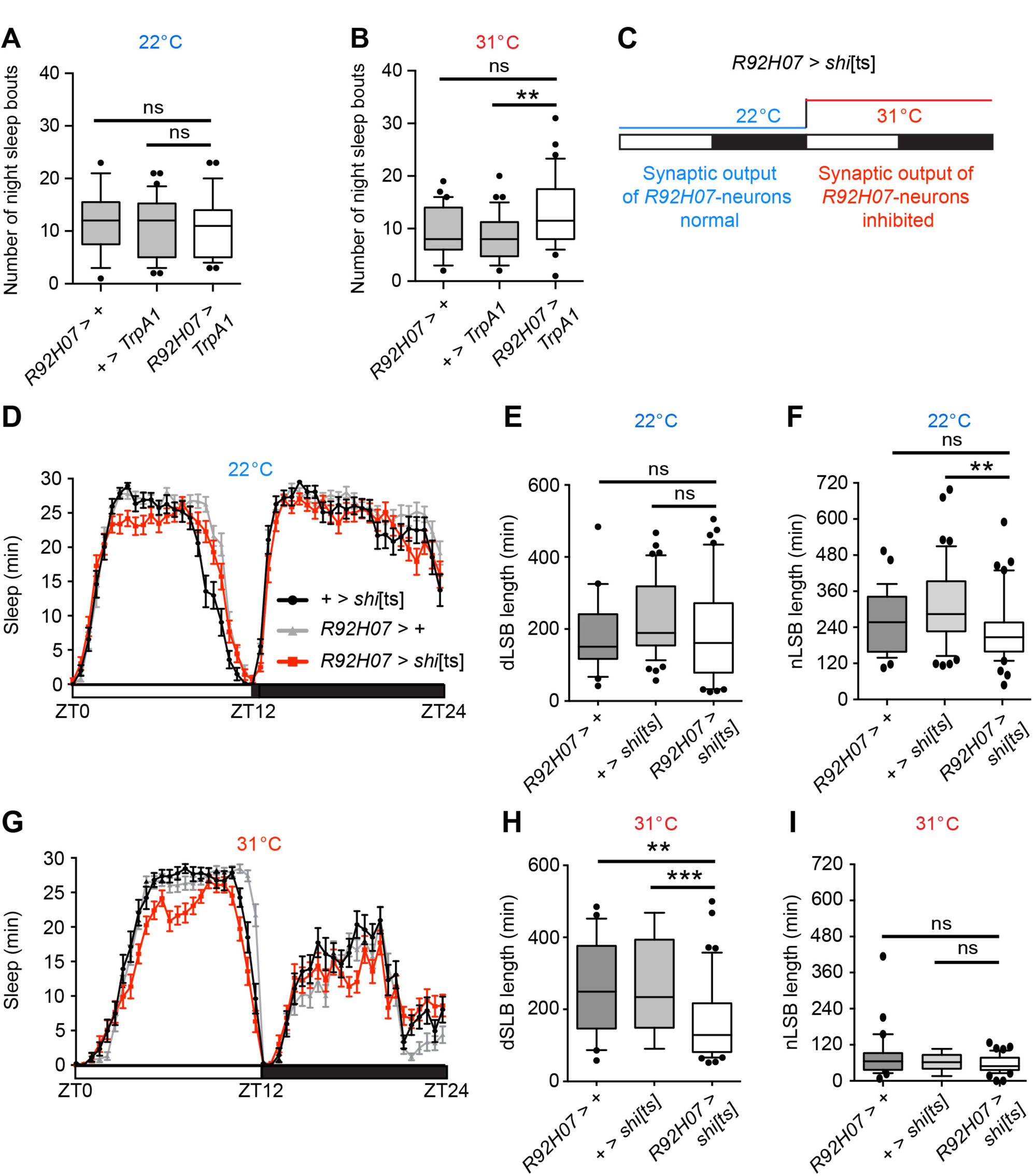
relating to Figure 3. Activation of *R92H07*-TuBu neurons does not cause paralysis. (A-B) Thermogenetic activation of *R*92*H*07-TuBu neurons does not alter the number of sleep bouts during the night. Sleep levels in flies expressing the TrpA1 thermo-sensory cation channel were measured on two consecutive days. On day 1 (A), ambient temperature was 22ºC, a non-activating temperature for TrpA1. On day 2 (B), ambient temperature was 31ºC, an activating temperature. Relative to both control lines, no change in sleep bout number during the night was apparent. Thus, when R92H07-TuBu neurons are activated, flies are still able to awaken and initiate movement, demonstrating that they are not paralysed. *R92H07* > +: n = 29; + > *TrpA1*: n = 34; *R92H07* > *TrpA1*: n = 36. (C) Paradigm for acute inhibition of *R92H07*-TuBu neurons at elevated temperatures using *shi*[ts]. (D-F) At the permissive temperature of 22ºC, the length of the longest day (dSLB) and night (nSLB) sleep bouts were unaffected by expression of *shi*[ts]. (G-I) At 31ºC, inhibition of synaptic release from *R92H07*-TuBu neurons results in a significant reduction in length of the dLSB but not the nLSB. *R92H07* > +: n = 45; + > *shi*[ts]: n = 29; *R92H07* > *shi*[ts]: n = 41. **p < 0.01, ***p < 0.001, ns – p > 0.05, Kruskal-Wallis test with Dunn’s post-hoc test.

